# Small mammals in biodiversity hotspot harbor viruses of epidemic potential

**DOI:** 10.1101/2024.09.10.612213

**Authors:** Yun Feng, Guopeng Kuang, Yuanfei Pan, Jing Wang, Weihong Yang, Wei-chen Wu, Hong Pan, Juan Wang, Xi Han, Lifen Yang, Gen-yang Xin, Yong-tao Shan, Qin-yu Gou, Xue Liu, Deyin Guo, Guodong Liang, Edward C. Holmes, Zihou Gao, Mang Shi

## Abstract

Metagenomic sequencing has transformed our understanding of viral diversity in wildlife and its threat to human health. Despite this progress, many studies have lacked systematic and ecologically informed sampling, which has left numerous potentially emergent viruses undiscovered, and the drivers of their ecology and evolution poorly understood. We conducted an extensive analysis of viruses in the lung, spleen, and gut of 1,688 animals from 38 mammalian species across 428 sites in Yunnan, China—a hotspot for zoonotic diseases. We identified 162 mammalian viral species, including 102 novel species and 24 posing potential risks to humans due to their relationships with known pathogens associated with serious diseases and their ability to cross major host species barriers. Our findings offer an in-depth view of virus organotropism, cross-species transmission, host sharing patterns, and the ecological factors influencing viral evolution, all of which are critical for anticipating and mitigating future zoonotic outbreaks.

## INTRODUCTION

Most pandemics, including that of coronavirus disease 2019 (COVID-19), are zoonotic in origin, initiated by the transmission of a microorganism from animals to humans [1–4]. Factors such as unchecked wildlife exploitation, climate change, and alterations in land use amplify human exposure to novel pathogens, increasing the risk of zoonotic disease emergence [2,3,5–7]. Although predicting exactly which virus may emerge next may not be possible [8–9], many viruses associated with human diseases had previously been identified in animal hosts [10–13]. As a consequence, the thorough surveillance of the viromes of animals that are closely linked to human populations is an important means to anticipate, mitigate, and prevent future zoonotic outbreaks [2,9,14–17].

Advances in metagenomic sequencing, particularly total RNA sequencing (i.e., metatranscriptomics) have deepened our understanding of viral characteristics in animal populations [18–26]. Recent research focusing on mammals closely associated with humans, especially the diverse mammalian order Rodentia and the sympatric small mammals of the orders Eulipotyphla (such as shrews) and Scandentia (i.e. tree shrews), has provided insights into potential zoonotic risks [23–26]. These animals inhabit varied terrestrial habitats with significant ecological overlap with humans [27] are the sources of major zoonotic pathogens such as the Lassa, Lymphocytic choriomeningitis, and Hantaan viruses, and play important roles in the natural cycles of vector-borne viruses like Crimean-Congo hemorrhagic fever and Tick-borne encephalitis viruses [28–30]. In addition, more structured metatranscriptomic surveys have enhanced our understanding of virome ecology and evolution, revealing both known viruses in new hosts and novel viral species that pose a threat to human populations [26, 31–34]. However, resource and logistical constraints often limit the scope of mammalian surveillance geographically, temporally, and numerically, resulting in a fragmented understanding of potentially emerging viruses and their ecological context.

Large-scale metagenomic studies often document virome compositions in host species at specific times and locations. This is particularly true of studies of small mammals such as rodents and shrews. In contrast, recent virome analyses of bats and mosquitoes that incorporated a broader ecological context not only identified high-profile pathogens, but also provided important insights into the extent and pattern of cross-species virus transmission as well as the determinants of virus biogeography [24,35], in turn providing a clearer picture of the drivers of disease emergence.

Situated in a 394,000 km² area and home to over half of China’s plant and vertebrate species, Yunnan province is a hotspot of global biodiversity [36]. Within this varied geographical landscape, we established extensive sampling sites across diverse environments to reveal the viromes of small mammals, including rodents, shrews and tree shrews, in doing so identifying viruses of potential zoonotic risk. Our data also addresses how both host and environmental factors impact virus richness, cross-species transmission, and genomic diversity.

## RESULTS

### 1. Sampling of Small Mammals in Yunnan

From 2021 to 2023, we conducted systematic animal sampling across Yunnanp province, China, capturing 1,688 mammals from 428 locations across all 16 prefectures and 96 counties (Fig. 1a and 1b, Table S1). The elevations of these locations ranged from 144 to 3,471 meters, and our sampling covered seven of the nine Köppen climate types in the province, excluding only two high-altitude types in the northwest (Fig. 1a and 1c, Fig. S1). Each captured animal was analyzed using the mitochondrial cox1 gene, with confirmed the presence of 38 mammalian species spread across 20 genera and 8 families (Fig. 1b). A rarefaction analysis indicated that the sampling strategy successfully captured a representative cross-section of the mammalian species diversity within the province (Fig. 1d). Rodents, predominantly from the genus *Rattus*, were the most frequently captured, with 1,540 individuals found at 419 sites. In contrast, shrews and tree shrews were relatively rare, with just 148 individuals from both groups captured across 49 sites. Notably, *Rattus* were widespread throughout the region, except in the warmest Cfa climate zone, where no samples were collected (Fig. 1c). Altitudinal analysis revealed species-specific distribution patterns, with *Apodemus chevrieri* found at higher elevations and *Rattus tanezumi* at lower elevations (Fig. S1).

**Figure 1.**
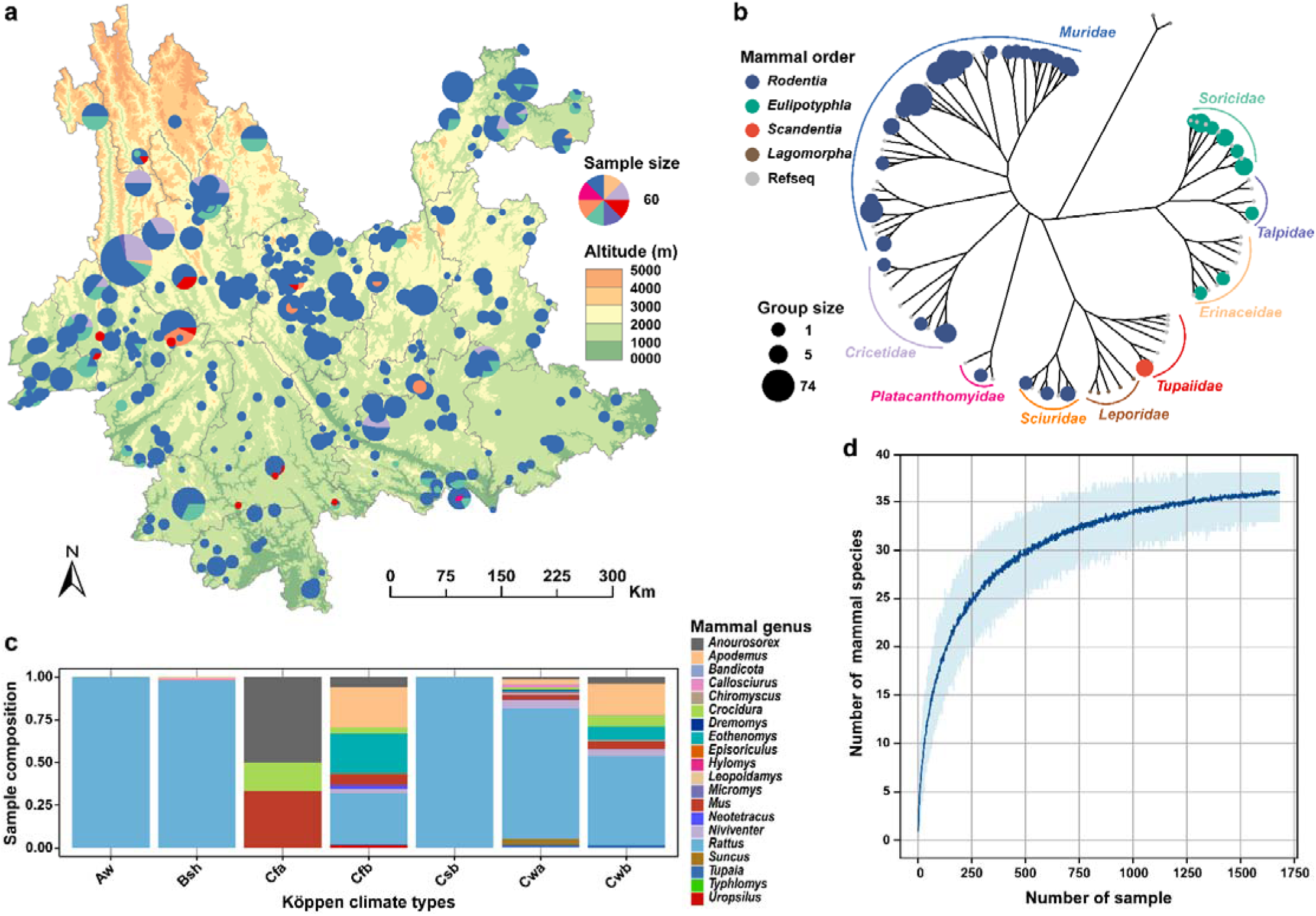
Distribution and diversity of small mammals in the study. **(a)** Map illustrating the sampling sites across Yunnan province, China, color-coded by digital elevation in ArcMap 10.8.1 (The basemap shapefile was sourced from the publicly accessible GADM dataset and ESDS). Pie charts at each site depict the mammalian family composition. **(b)** Phylogenetic tree based on the cox1 gene of mammals analyzed in this study. Circle sizes reflect number of sample groups and colors represent mammalian orders. **(c)** Composition of mammalian genera across different Köppen climate types, with segments color-coded to reflect the relative abundance of each genus. **(d)** Rarefaction curve demonstrating the relationship between the number of samples collected and the mammalian species identified.

We strategically pooled 7-8 individuals from the same species and location into 207 sample groups for metatranscriptomic sequencing, guided by mitochondrial sequence data (Table S2). For species with fewer than 7 individuals from a specific region, samples were merged across broader areas, creating an additional 18 groups (Table S2; see Materials and Methods for details). Consequently, a total of 225 sample groups were organized. Metatranscriptomic libraries were prepared from organ tissues (gut, lung, and spleen) for each group. A total of 646 libraries produced 21.99 billion clean non-rRNA reads, averaging 34.04 million reads per library (Table S2).

### 2. Composition and diversity of mammalian virome

Our analysis revealed an extensive diversity of viruses across host species. We focused on viruses associated with mammalian infections; specifically, those phylogenetically related to known (i) vertebrate-specific viruses and (ii) to vector-borne viruses that are associated both mammals and arthropods (Fig. 2 and S2). Accordingly, our study classified 5,350 viral contigs into 162 mammalian virus species across 23 families, including 116 RNA virus species from 16 families, 41 DNA virus species from 7 families, and 5 reverse transcribing virus species from the *Retroviridae* and *Hepadnaviridae* (Fig. 2b). Notably, 103 species from 19 families were newly identified species according to the ICTV species demarcation criteria (Table S3). Of the 646 libraries analyzed, 414 contained mammalian viruses, with viral RNA constituting between 1.08L×L10^−6^ % and 15.43% of the total clean non-rRNA reads per library. Among these, the median number of mammalian viruses detected in each library was three, with an interquartile range of four, and a maximum of 16 viral species (Fig. 2a). In total, 204 of 225 sampled groups were discovered at least one mammalian virus.

**Figure 2.**
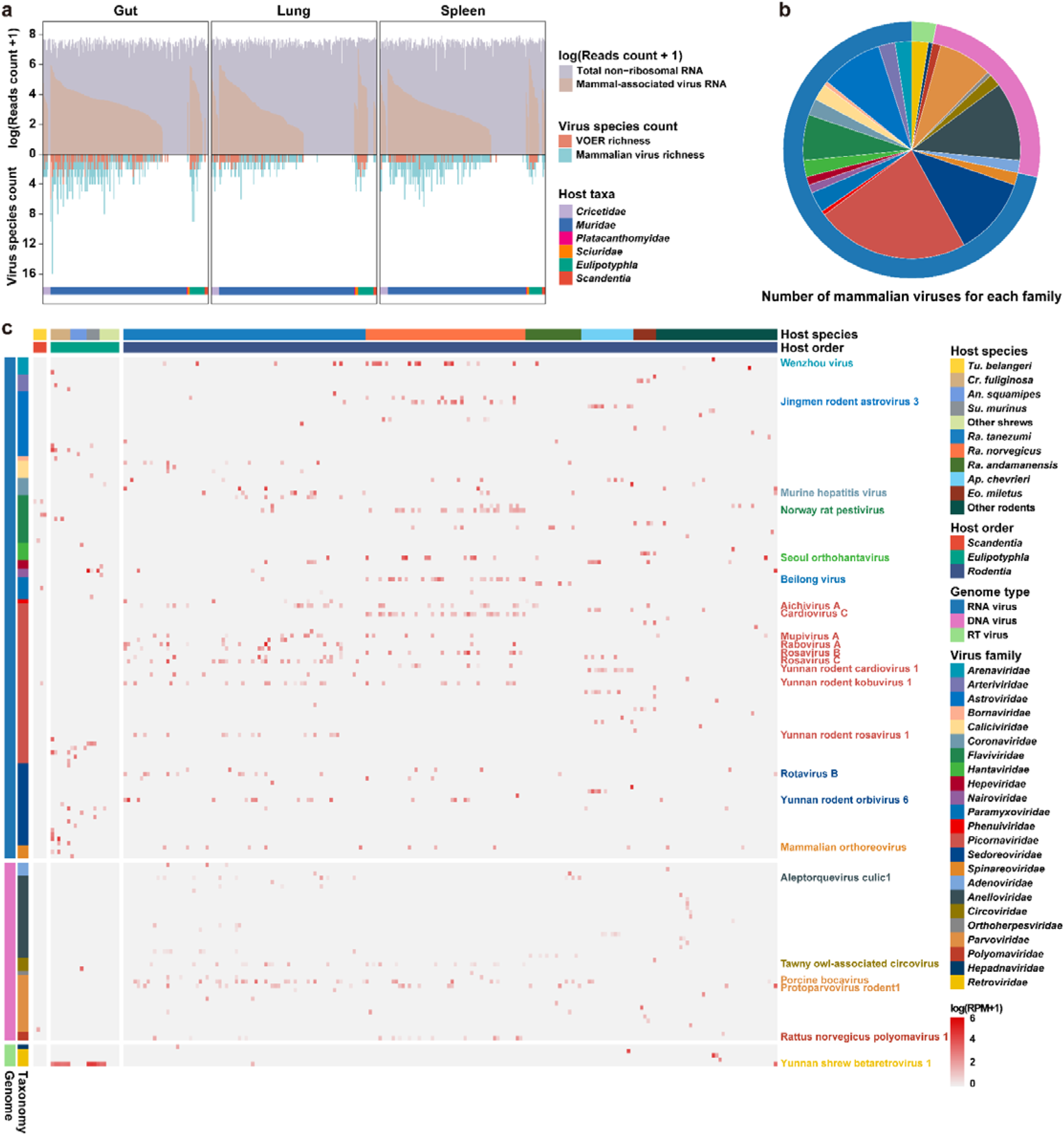
Overview of the diversity and prevalence of mammalian viruses in this study. **(a)** Total read depth, viral read abundance, and species richness of viruses from each library, segmented by organ type: gut, lung, and spleen. **(b)** Pie chart showing the distribution of mammalian virus species identified within each virus family. **(c)** Heatmap of the distribution and relative abundance of mammalian viruses, quantifying viral abundance in each sample group by mapped reads per million non-rRNA reads (RPM). Host species and orders are labeled at the top, color-coded to match their respective categories. Viral species from 23 families are displayed, with each family distinctly colored; names are provided for only those viruses identified in more than 10 groups to emphasize the most prevalent species.

The diversity and prevalence of viral families varied significantly. *Picornaviridae* was the most diverse, with 37 species identified, followed by the *Sedoreoviridae* and *Anelloviridae*, each with 19 species. In contrast, the *Bornaviridae* and *Phenuiviridae* were each represented by only one identified species (Fig. 2b). Commonly detected families included the *Picornaviridae*, *Sedoreoviridae*, *Flaviviridae*, and *Parvoviridae*, found in at least 10 sample groups each, whereas families such as the *Bornaviridae*, *Hepadnaviridae*, and *Orthoherpesviridae* appeared more sporadically (Fig. 2c).

### 3. Identification of viruses of emergence risk

Among the 162 mammalian viral species identified here, 24 were designated as ‘viruses of emergence risk’ (VOER) based on their close phylogenetic relationships to known human pathogens (Table S4) and/or their increased risk of cross-species transmission (Fig. 1a). Indeed, 20 of these VOERs were closely related to known human pathogens (Fig. 3b, Table S4), highlighting their epidemic potential.

**Figure 3.**
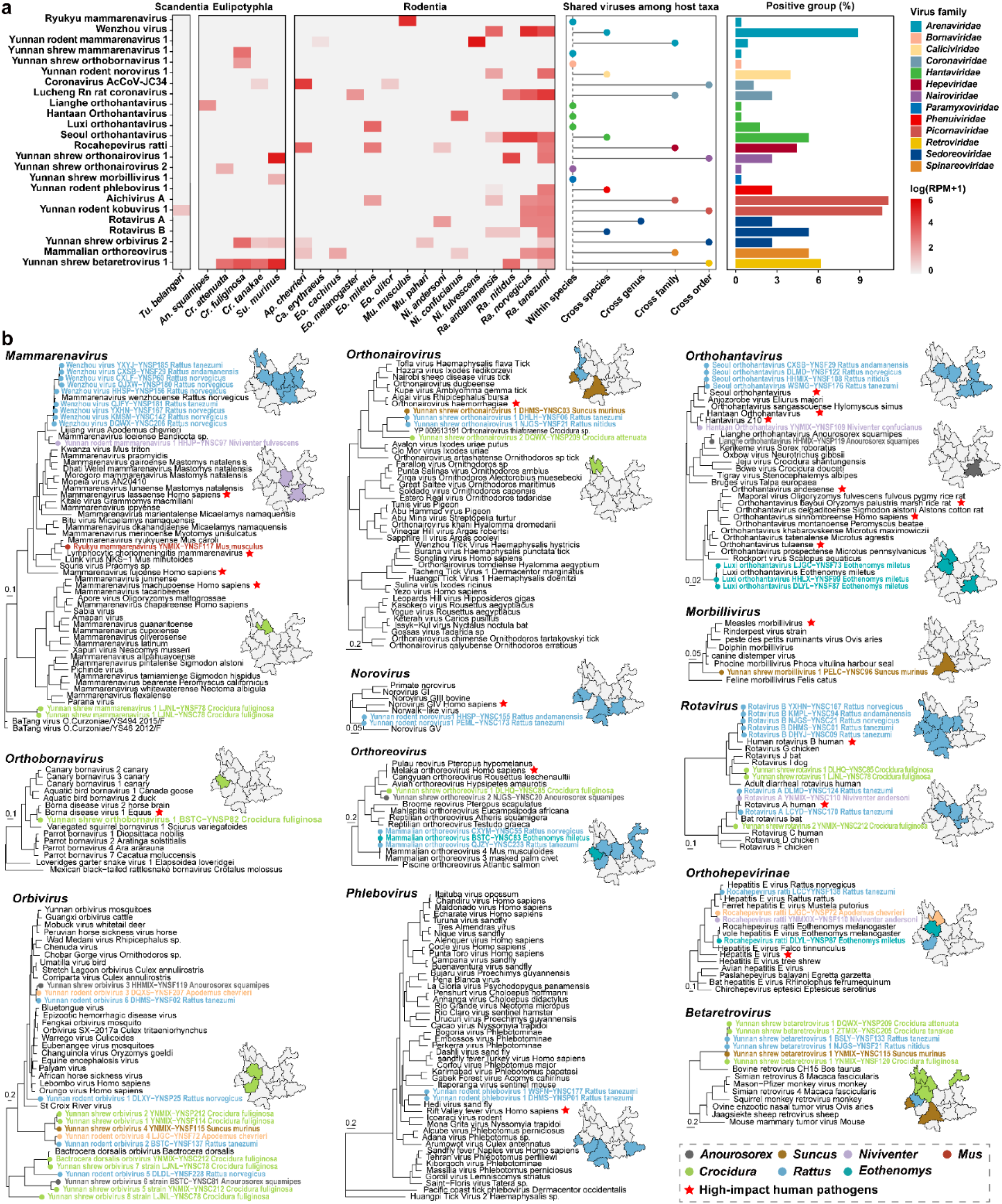
Phylogenetic relationships and epidemiological characteristics of viruses of emergence risk (VOER). **(a)** Mammalian host distribution, extent of virus sharing among hosts, and prevalence of viruses. The heatmap illustrates the abundance of each virus within different host species, the middle bar chart depicts how viruses are shared among host taxa, while the right bar chart shows the prevalence of each viral species. (**b)** Phylogenetic relationships and geographical distribution of VOERs. Phylogenetic trees for each established human-infecting virus genus were estimated using a maximum likelihood method, based on conserved viral proteins (RdRP for RNA virus and Reverse transcriptase for the *Retroviridae*). Trees were midpoint rooted, with branch lengths representing the number of amino acid substitutions per site. Dots on the trees, colored according to host genera as indicated in the legend, represent viral species identified in this study. Previously identified human pathogens in each genus are marked with red pentagrams. The accompanying maps, with colors matching the strain names in the trees, illustrate the geographic distribution patterns of these viruses.

In the context of viral groups often associated with hemorrhagic fever or encephalitis, seven species of arenavirus and hantavirus were identified, including three that are newly discovered. Notably, *Yunnan shrew orthobornavirus 1*, exhibited high genetic similarity to classic *Borna disease virus 1* (87.6% amino acid identity the large protein) and *2* (88.5% identity), and was also related to *Variegated squirrel bornavirus 1* (74.8%) identity, known to cause fatal human encephalitis [37] (Fig. 3b, Table S4).

In addition, we identified several new arthropod-borne viruses (i.e., arboviruses): one phlebovirus (*Yunnan rodent phlebovirus 1*) and two orthonairoviruses (*Yunnan shrew orthonairovirus 1* and *2*) that were closely related to *Rift Valley fever virus* and *Crimean-Congo hemorrhagic fever virus*, respectively, both of which can lead to lethal hemorrhagic fever, such that the newly discovered virus may also pose a threat to humans (Fig. 3b, Table S4). Moreover, 15 species of orbivirus, again likely arboviruses, were discovered.

In the case of potential respiratory infections, we identified *Yunnan shrew morbillivirus 1* that exhibited 66.3% sequence identity to *Measles morbillivirus*. Additionally, *rotavirus A* and *B*, *Yunnan rodent norovirus 1* and *Aichivirus A* were flagged as potential disease agents related to these enteric human viruses (Fig. 3b).

Several viruses were classified as VOER due to their capacity for transmission among genetically distant mammalian species, demonstrating that they may represent host generalists (Fig. 3a). For instance, Lucheng Rn rat coronavirus and coronavirus ArcCoV-JC34 were identified in more than two different mammalian families, while Yunnan shrew betaretrovirus 1 and Yunnan shrew orbivirus 2 were present in different mammalian orders.

Interestingly, we also discovered viruses that occupy unique phylogenetic positions in the mammalian-associated virus lineages. For example, seven novel virus species carried by shrews fell as basal lineages to the genus *Orbivirus*, order *Reovriales*; *Yunnan shrew mammarenavirus 1* was phylogenetically positioned between newly identified *BaTang virus* and the classic Old World and New World mammarenaviruses; and *Yunnan shrew morbillivirus 1* was basal to the entire genus *Morbillivirus* (Fig. 3b). Interestingly, these viruses were predominantly found in shrews from the genera *Anourosorex*, *Crocidura*, and *Suncus*, suggesting that these taxa harbor diverse viromes.

### 4. Organotropism of mammalian viruses

Of the 225 groups sampled, 201 had complete set of all three organs (gut, lung, and spleen), leading to 603 libraries that were used for organotropism analysis. Significant differences in virome composition and abundance were observed across these organs (adonis test, R² = 0.28, p < 0.01; Fig. 4a and S3). Additionally, an analysis of virus detection frequencies across different organs revealed significant variation in viral distribution (Chi-squared test, X² = 667.88, p < 0.05). Based on relative abundance, distinct organotropisms were noted for many viral families: *Hantaviridae* and *Nairoviridae* were primarily found in lungs, *Phenuiviridae*, *Arteriviridae* and *Circoviridae* in spleen, and *Picornaviridae*, *Caliciviridae*, and *Coronaviridae* in the gut (Fig. 4b). Conversely, families such as the *Retroviridae*, *Arenaviridae*, *Hepeviridae*, *Parvoviridae* and *Sedoreoviridae* appeared in all three organs, suggesting systemic infections (Fig. 4b). Similar patterns of organ-specific and systemic infections were also noted at the viral species level (Fig. 4c, 4d and S3). Interestingly, some virus species, such as Protoparvovirus rodent 1, Yunnan shrew betaretrovirus 1, and Yunnan shrew hepatovirus 1, despite showing systemic infection, exhibited uneven viral abundances across different organs, indicating distinct organ preferences (Fig. 4d).

**Figure 4.**
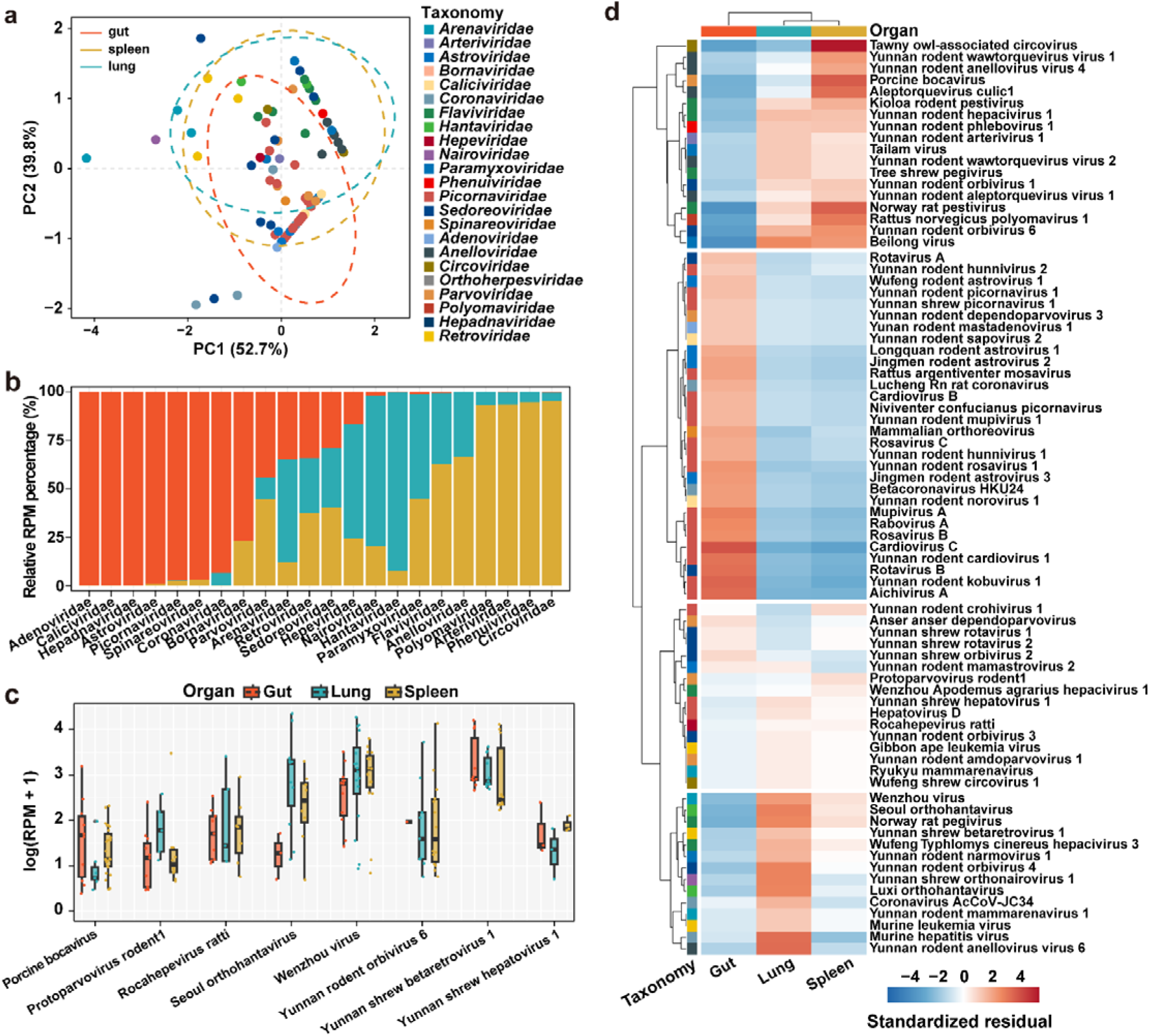
Organ distribution patterns of mammalian viruses. **(a)** Principal component analysis (PCA) based on the median viral abundance (RPM) of each viral species (colored by family), highlighting virome composition across the gut, spleen, and lung. **(b)** Relative total viral abundance (RPM) for various viral families in gut (orange), lung (turquoise), and spleen (golden yellow). **(c)** Comparison of the abundance of viruses detected in at least three libraries for three organ types, with colors keys shown to the legend. **(d)** Heatmap illustrating organotropism of viruses, determined by a chi-squared test analyzing distribution frequencies of virus species across gut, lung, and spleen.

### 5. Virome composition and transmission dynamics

To analyze virus composition and transmission among different mammalian hosts, we focused on groups with complete data for three specific organs and excluded those with fewer than seven individuals or fewer than three sampled groups per genus. Consequently, from the orders Eulipotyphla (23 individuals, 3 groups), Scandentia (91 individuals, 12 groups), and Rodentia (1255 individuals, 159 groups), we identified 6, 29, and 106 viruses, respectively (Fig. S4). Virus richness and distribution varied significantly (Fig. 5a): Rodentia (i.e. rodents) primarily hosted members of the *Picornaviridae*; Eulipotyphla (i.e., shrews) mainly harbored *Sedoreoviridae*; and Scandentia (i.e. tree shrews) commonly exhibited *Flaviviridae*.

**Figure 5.**
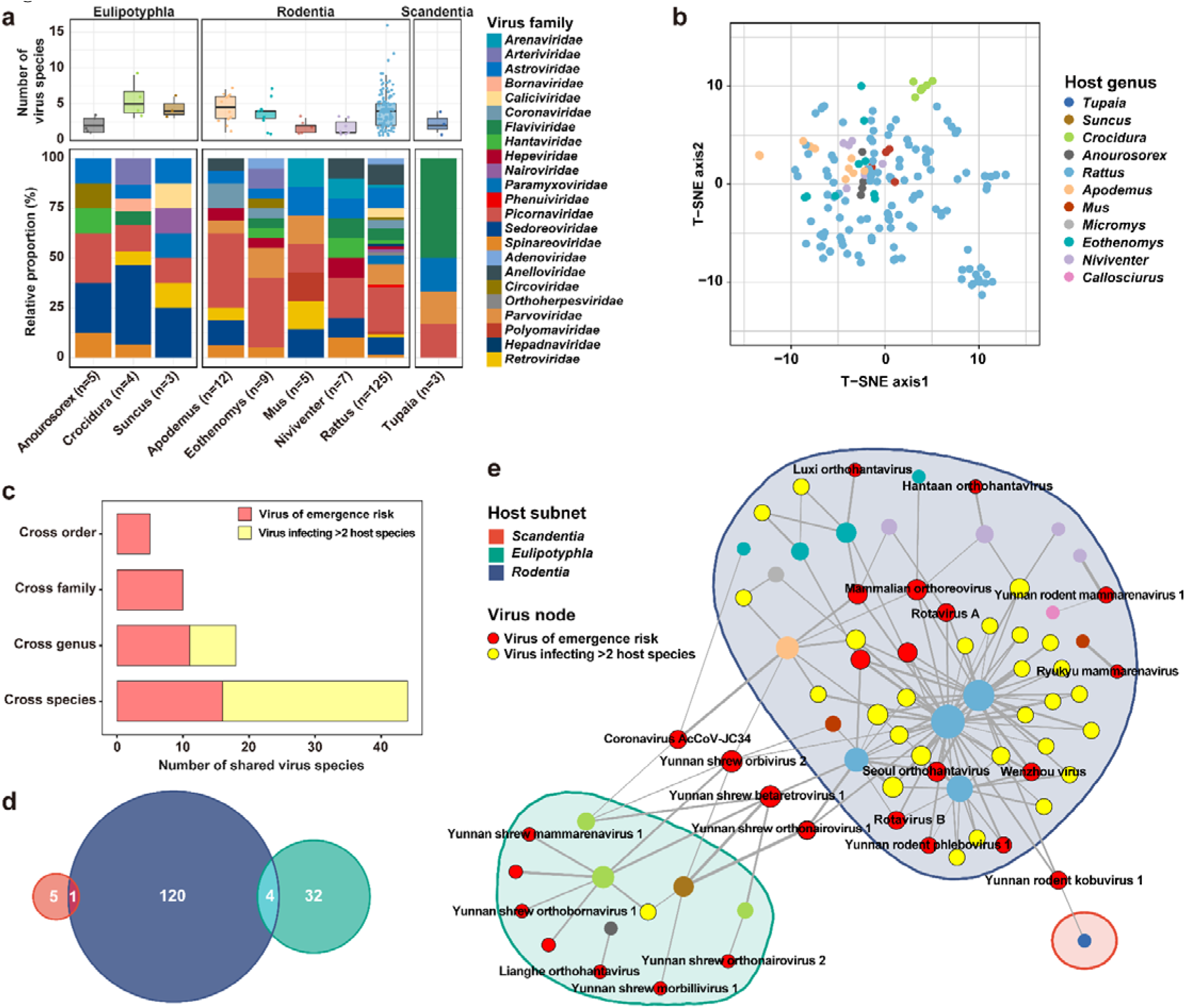
Viral composition and transmission among mammalian hosts. **(a)** Number of viruses per sample group and their distribution across mammalian taxa. **(b)** t-SNE ordination illustrating the distinct virome compositions of mammalian genera, with each point representing a group colored according to the host genus as indicated in the legend. **(c)** Number of viruses shared across species, genera, families, and orders, highlighting extensive virological sharing among different mammalian hosts. **(d)** Venn diagram depicting specific viruses shared among the mammalian orders of Scandentia (red-orange), Eulipotyphla (teal), and Rodentia (dark blue). **(e)** Virus sharing network: nodes represent hosts or virus species, colored by host genera and virus types. Line thickness between nodes indicates the relative abundance of viruses, with the network divided into three subnets based on host order. External nodes emphasize viruses with potential for cross-order transmission.

A t-distributed stochastic neighbor embedding (t-SNE) analysis visualized distinct virome compositions across host taxa (Fig. 5b and S4). PERMANOVA tests on viral genera with at least five sampled groups confirmed significant differences in virome compositions among mammalian genera (R² = 0.09, p < 0.001) and species within the same genus (p < 0.001), although the latter showed a lower variance (R² = 0.07), indicating less pronounced differences at the intra-genus level.

We also identified several instances of virus sharing among different host taxa, indicative of potential cross-species transmission. A total of 44 viral species were detected in at least two host species, with 18 viruses found across different genera, 10 across different families, and 5 across different orders (Fig. 5c). Scandentia and Rodentia shared one viral species, while Eulipotyphla and Rodentia shared five (Fig. 5d). No viruses were shared across all three orders. Notably, all 10 viruses shared across mammal families, including five shared between orders, were identified as VOER (Fig. 5e).

### 6. Determinants of viral richness, composition and intra-specific genomic diversity

A total of 94 sample groups with suitable sample size and range were selected for ecological comparisons to explore the environmental and host factors influencing viral species richness, composition, and genomic sequence diversity. Generalized linear models (GLMs) were utilized for these analyses. We selected the best model structures (ΔAIC <2) by evaluating all combinations of variables based on the Akaike Information Criterion (AIC) (Fig. S5). This analysis identified that the key determinants varied among different metrics of viral diversity (Fig. 6a).

**Figure 6.**
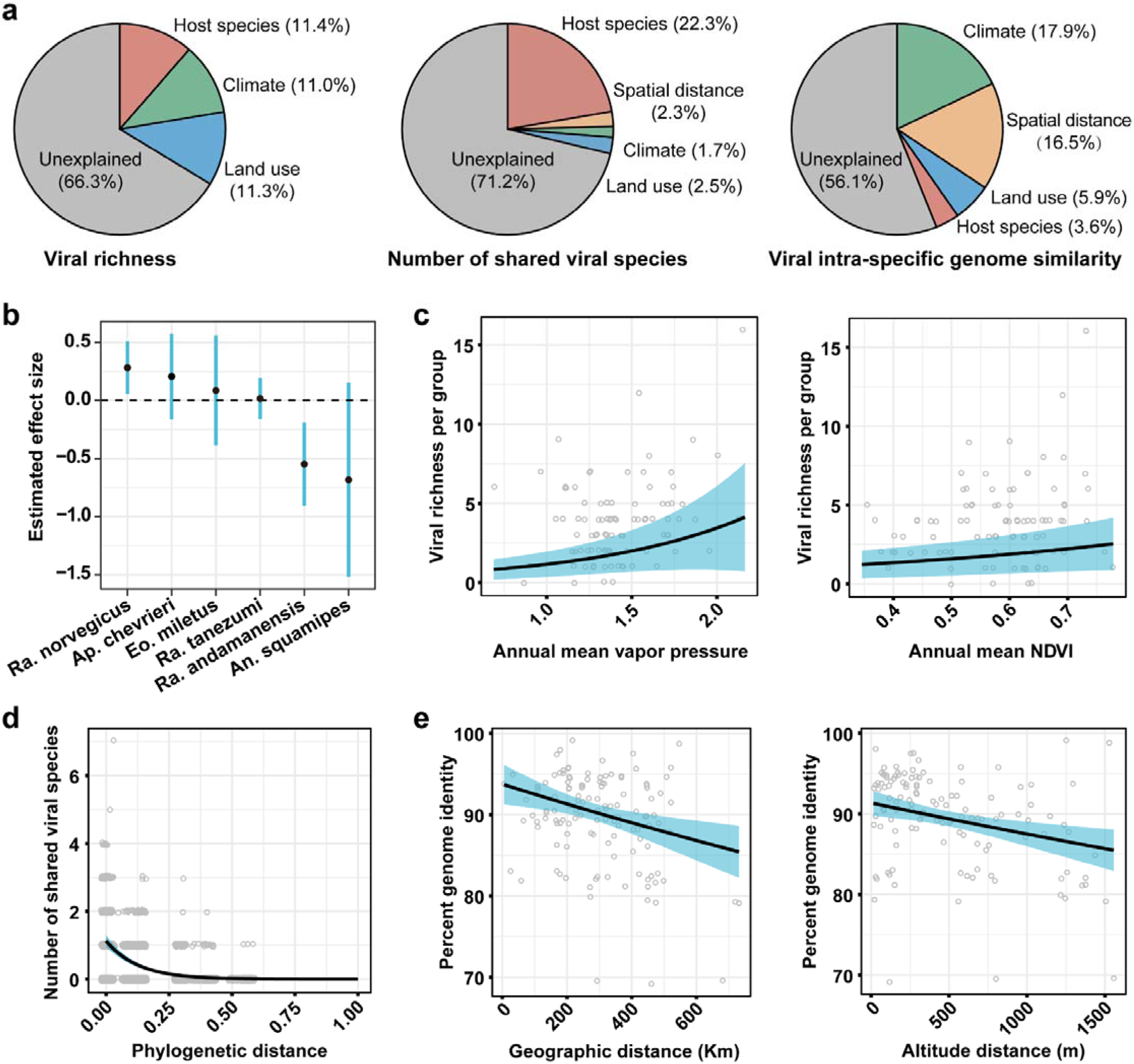
Key determinants of viral species richness, composition, and intra-specific genomic diversity. **(a)** Analysis of the relative contribution of host species, climate, and land-use characteristics to viral richness in each sampled group, quantified by the explained deviance in the best model structures (ΔAIC <2) using generalized linear models (GLMs). **(b)** Model estimates showing the impact of mammalian species on viral richness per sampled group, presented with estimated values andL95% confidence intervals (CI). **(c)** Relationship between annual mean vapor pressure (a climate variable) and annual mean NDVI (a land-use variable) with viral richness in each sample group. **(d)** Relationship between the number of shared viral species and host phylogenetic distance (patristic distance based on the cox1 phylogeny) among pairs of sample groups. **(e)** Impact of spatial distance (horizontal geographical distance and vertical altitudinal distances) on intra-specific viral genome similarity.

Variations in viral richness were influenced by host species (11.4%), climate (11.0%), and land-use variables (11.3%), with 66.3% of the total deviance remaining unexplained (Fig. 6a). Indeed, the distribution of viral richness across mammal species was uneven (Kruskal-Wallis H test, p < 0.05; Fig. 6b). Individual climatic variables, such as annual mean vapor pressure, and land-use variables like vegetation density (normalized difference vegetation index, NDVI), positively influenced viral richness (Fig. 6c). Furthermore, separate analyses for the two most abundant rodent species – *Rattus tanezumi* (n = 41) and *Rattus norvegicus* (n = 20) – revealed that climatic conditions CPC1 (see Methods for details) and NDVI were the primary determinants, explaining 48.5% of the total variance in viral richness for *Rattus tanezumi*. In contrast, for *Rattus norvegicus*, viral richness was chiefly linked to human population density, accounting for 33.9% of the total variance (Fig. S5).

In contrast, viral composition, measured as shared viral species between pairs of sampled groups, were most significantly influenced by host phylogenetic distance, which explained 22.3% of the deviance (Fig. 6a). After adjusting for other covariates such as climate differences, land-use variables, and spatial distance, the number of shared viral species notably decreased as phylogenetic distance between host groups increased (Fig. 6d). Conversely, the number of shared viral species was slightly negatively affected by differences in climatic variables (annual mean climatic water deficit), land-use characteristics (cropland presence), and mammal richness (Fig. S5).

Finally, viral intra-specific genetic diversity (genome similarity) was primarily influenced by climate (17.8%) and spatial distance (16.6%), with the latter being a significant contributor unlike in the analysis of viral richness and composition. Indeed, viral genome similarity significantly decreased with increasing spatial distance, encompassing both horizontal geographical and vertical altitudinal dimensions (Fig. 6e and S5). In contrast, the association with host phylogenetic distance was relatively weak.

## DISCUSSION

Our study identified a number of viruses in rodents, shrews and tree shrews that may also have the capacity to emerge in humans, including both newly identified and previously known viruses associated with such syndromes as hemorrhagic fever, fever and encephalitis. This highlights the high risk of disease emergence at human-animal interfaces in Yunnan province, where we found a greater variety of such zoonotic viruses than previously reported [26,31–34]. The well-established correlation between viral diversity and zoonotic risk with reservoir richness, population size, and density [21,38–39] supports these findings. Indeed, the viral diversity observed here can in large part be attributed to Yunnan’s diverse geographical landscapes and its rich biodiversity. This region is globally recognized for its dense distribution of significant wildlife taxa and is a critical terrestrial biodiversity area [36]. Additionally, our study benefits from the first broad and systematic sampling effort in this region, which covered a wide diversity of hosts and densely sampled areas. This comprehensive approach has deepened our understanding of which wildlife species and viruses pose significant risks to humans and clarified the contexts in which these viruses are likely to emerge.

Our discovery of many potentially zoonotic viruses can also be attributed to our extensive and separate examination of three organs (namely lung, gut, and spleen), each representing critical systems—respiratory, digestive, and circulatory—that are potential routes for infection through animal aerosols, excretions, and arthropod bites. This multi-organ approach has broadened our understanding of the range of pathogens posing threats to human health, overcoming the limitations of previous studies that targeted only specific tissues or swabs and likely overlooked significant portions of the virome [32–33]. Indeed, our data indicate that each organ might harbor unique viral pathogens. For instance, vector-borne pathogens were primarily found in the spleen. Interestingly, we detected a circovirus that was significantly enriched in the blood, differing from its typical manifestation in other mammals [40–41]. Additionally, our research identified viruses with multi-organ distributions, indicative of systemic infections, including several divergent members of the genus *Orbivirus*. The detection of these systemic infections highlights the critical role of rodents and shrews as likely principal hosts for these viruses, emphasizing their significance in the transmission of diseases.

Our study uncovered a large and intricate network of virus-host transmission among small mammals in Yunnan, shedding light on potential zoonotic threats. On one hand, we found that cross-species virus transmission occurs frequently, with 44 (27.16%) virus species carried by at least two host species. Conversely, our results also revealed strong host restrictions on cross-species transmission, suggesting that the greater the genetic distance between hosts, the less likely virus transmission will occur. This pattern was underscored by host genetic distance being the single dominant factor influencing viral composition, consistent with previous studies on other organisms [14,24,35,42–43]. This explains why despite the frequent occurrence of cross-species transmission, spillover to humans resulting in zoonotic diseases is typically rare [17,21,44]. Despite this, some viruses demonstrate a notable capacity to infect genetically distant hosts: we identified 10 viral species present in at least three host species from different mammalian families, and four viral species across different mammalian orders. These viruses, which we denote as VOER, merit particular attention due to their broad host range [14,23]. For instance, viruses capable of overcoming major host barriers, such as SARS-CoV-2 – which uses the conserved ACE2 receptor to infect a wide range of mammalian species other than humans – highlight the great potential for zoonotic transmission [4,45–46]. Thus, the identification and close monitoring of such viruses are crucial for preventing zoonotic events.

Our extensive sampling across wide environmental gradients provided a robust data set to investigate the ecological factors influencing viral diversity among mammalian wildlife. We found that viral richness, composition, and intra-specific genetic diversity were driven by a combination of host species, climatic conditions, and land-use variables. Specifically, viral richness was significantly influenced by host species diversity, annual mean vapor pressure, and vegetation density, accounting for a considerable portion of the observed variance. While it is known that viral diversity may be affected by host, climate, and land-use factors, such effects have seldom been quantitatively assessed. Our results provided a quantitative test of these effects, aligning with previous studies that highlight the role of environmental and ecological variables in shaping viral diversity. For instance, a previous study demonstrated that host taxonomy and land-use types (mountainous versus agricultural) were critical determinants of viral richness in bats, rodents, and shrews [26]. However, in contrast to that study, we observed a greater frequency of viruses in rodents than shrews. Although our sampling was strongly biased toward rodents, diversity-hotspot hosts may vary spatially.

Our findings distinguish between inter-specific diversity (measured by the number of virus species shared among samples) and intra-specific diversity (measured by viral genomic sequence similarity among samples), showing they are influenced by different factors. Inter-specific viral sharing is primarily driven by the phylogenetic distance between hosts, which significantly affects virus species sharing, while climate and spatial distance have less or negligible influence. However, this analysis treats each virus species as a whole without considering sequence variation within species. When we examine diversity at a finer evolutionary scale, focusing on genomic similarities within each viral species, both differences in climate and spatial distances—especially along vertical elevational gradients—emerge as the dominant factors. This suggests that climatic and spatial factors do impact virus diversity, but this influence is more pronounced on recent evolutionary scales. Despite this, overcoming spatial barriers is generally easier for viruses than overcoming host species barriers, with the latter clearly a major determinant of virus evolution [47–48].

### Limitations of the study

There were several limitations to our study. While the reliance on pooled samples for metatranscriptomic sequencing is practical for broad surveys, this approach might obscure intra-species viral diversity and the detection of low-abundance viruses. Additionally, there is potential bias introduced by the uneven sampling across different climate zones and elevations. Although we attempted to capture a representative cross-section of mammalian species, certain areas, particularly the warmest and highest-altitude zones, were underrepresented. Another limitation is the focus on a specific geographic region, which may limit the generalizability of our findings to other areas with different ecological and climatic conditions. Despite these limitations, we were able to integrate ecological, climatic, and host-related factors to elucidate the factors that shape viral diversity. This highlights the need for more extensive and geographically diverse studies to fully capture the global patterns of viral evolution and transmission.

## MATERIALS AND METHODS

### 1. Study design and sample collection

This study characterized the viromes of a broad spectrum of small mammalian species to investigate the diversity of viruses they harbor and the ecological factors that shape virome composition. To ensure a representative sample across diverse ecological settings, 428 sampling sites (villages) across 96 counties in all 16 prefectures of Yunnan province, China, were systematically surveyed from 2021 to 2023. The number of sites varied from 1 to 22 per county, and from 4 to 108 per prefecture. These sites included both urban and rural areas, covered 7 of the 9 Köppen climate types (https://www.britannica.com/science/Koppen-climate-classification), and spanned altitudes from 144 to 3471 meters (Fig. 1a and Fig. S1, Supplementary Table 1 and 2). In total, 1540 rodents, 125 shrews, and 23 tree shrews were captured (Supplementary Table 3). Rarefaction analysis confirmed that the captured small mammals adequately represented true species diversity (Fig. 1d).

Rodents, shrews and tree shrews were captured using snap-traps or cage-traps. Following capture, geographic coordinates and altitudes were recorded. Animals were then euthanized and dissected. Gut (including feces), spleen, and lung tissues were harvested and immediately preserved in dry ice or liquid nitrogen, and subsequently stored at −80°C. All protocols for sample collection and processing were reviewed and approved by the Ethics Committee of the Yunnan Institute of Endemic Diseases Control and Prevention and Sun Yat-sen University (SYSU-IACUC-MED-2021-B0123).

Mammalian species were initially identified by experienced field biologists based on morphological characteristics. This preliminary identification was confirmed by sequencing and analyzing cytochrome c oxidase (COI) gene for each specimen [49]. For each library, mammalian species confirmation was achieved using *de novo* assembled contigs of COI genes. The final clean cox1 contigs were submitted to the BOLD online system [50] for species identification. A phylogenetic tree incorporating all full-length COI sequences was estimated using PHYML 3.0 [51], employing the GTR+Γ nucleotide substitution model and the Subtree Pruning and Regrafting (SPR) branch-swapping algorithm.

### 2. Sample groups

Before RNA extraction, individual animals were first organized into sample groups. Typically, each group comprised 7-8 animals of the same species and location. In cases where fewer than 7 animals were available from the same conditions, animals sampled from wider geographic areas (first from the same prefecture, then from the entire province) were combined into groups. Of these, 207 groups comprised 7-8 animals, while 18 groups included fewer than 7 animals. In total, this approach resulted in 225 sample groups for the study (Supplementary Table 2).

### 3. RNA extraction, library construction and sequencing

Tissue samples from each organ (gut, lung, and spleen) and each animal were individually homogenized in 600 µl of MEM solution (GIBCO). The homogenates were then pooled by organ type in equal volumes to create 200 µl pools for each sample group. As a result, each sample group had three distinct pools corresponding to the gut, lung, and spleen. Total RNA was extracted and purified from each pool using the RNeasy Plus Universal Mini Kit (Qiagen, Germany). RNA libraries were then constructed using the Zymo-Seq RiboFree™ Total RNA Library Kit (No. R3003), according to the manufacturer’s instructions. These libraries were sequenced using paired-end (150 bp reads) on the Illumina NovaSeq 6000 sequencing platform.

### 4. Identification and confirmation of mammalian viruses

For the raw sequencing reads from each library, adapters were removed and initial quality control was conducted using the pipeline implemented in the bbduk.sh program (https://sourceforge.net/projects/bbmap/). The parameters for adapter removal included ktrim=r, k=23, mink=11, hdist=1, tpe, tbo. Quality control settings were maq=10, qtrim=r, trimq=10, ftl=5, minlen=90. Reads with extensive non-complex regions were excluded (parameters: entropy=0.5, entropywindow=50, entropyk=5). Duplicate reads were filtered out using cd-hit-dup under default settings [52]. rRNA reads were then removed by mapping the processed reads against the SILVA rRNA database (Release 138.1) using Bowtie2 (version 2.3.5.1) in the ‘--local’ mode [53]. The remaining high-quality, non-rRNA reads underwent *de novo* assembly with MEGAHIT (version 1.2.8) using default parameters [54]. The assembled contigs were then analyzed using DIAMOND BLASTx against the NCBI non-redundant protein database [55] with an e-value threshold of 1×10^−5^. Taxonomic classifications were assigned by correlating the top BLAST hit accession numbers to NCBI taxids, extracting those identified under the kingdom ‘Viruses’ for subsequent analyses.

Viral contigs shorter than 600 bp were excluded to ensure quality, and the remaining overlapping unassembled contigs were merged to form extended viral sequences using the SeqMan program implemented in the Lasergene software package (version 7.1, DNAstar) [56]. To verify genome integrity for viral families such as the *Retroviridae, Hepadnaviridae,* and *Bornaviridae* that may integrate into mammalian host genomes, open reading frames (ORFs) were identified using ORFfinder (https://www.ncbi.nlm.nih.gov/orffinder/) and sequences were classified via the online BLASTp program (https://blast.ncbi.nlm.nih.gov/Blast.cgi). The abundance of these viral contigs was estimated by mapping reads back to the assembled genome with Bowtie2 (version 2.5.2) using ‘--end-to-end’ and ‘--very-fast’ settings. Alignments were sorted and indexed with SAMtools (version 1.18) and visualized with Geneious Prime (version 2020.2.4) [57–58]. To further verify these findings and eliminate false positives, contigs were cross-referenced against the non-redundant nucleotide database using online BLASTn to exclude sequences related to the host genome, endogenous viral elements, and artificial vectors.

Each viral contig was classified at the species level based on the species demarcation criteria established by the ICTV for the viral genus in question [59]. For genera lacking explicit species demarcation criteria, a 90% amino acid identity threshold for the RdRP or replicase protein was applied (Supplementary Table 3). The classification of viral contigs was further validated through comparisons of amino acid sequences from conserved genes, which included the RdRp for RNA viruses, pol for the *Retroviridae*, the major capsid protein for the *Orthoherpesviridae*, LTAg for the *Polyomaviridae*, ORF1 protein for the *Anelloviridae*, NS1 for the *Parvoviridae*, and DNA polymerase for other DNA viruses. Alignment of draft virus sequences and corresponding reference sequences from GenBank were performed using MAFFT (version 7.48) [60], with ambiguously aligned regions removed using TrimAl^61^. Phylogenetic trees were estimated using the maximum likelihood (ML) method implemented in PHYML 3.0 [51], employing the LG model of amino acid substitution and SPR branch-swapping. Only viral contigs that phylogenetically clustered with recognized mammalian-infecting viruses were considered mammalian viruses and retained for further analysis.

### 5. Viral RNA quantification

To quantify the amount of viral RNA present in our samples, we constructed reference sequences for all sequence variants of each viral species. Viral abundance was measured in each library by counting the number of viral reads per million of the non-rRNA reads (RPM). Reads were mapped to these reference genomes using Bowtie2 with ‘--end-to-end’ and ‘--very-fast’ settings, ensuring accurate alignment and quantification. To mitigate potential false positives from index-hopping, viral reads were only considered valid if they accounted for more than 0.1% of the highest read count within the same sequencing lane. Additionally, data characterized by low abundance (RPM < 1) or insufficient genome coverage (< 300 bp) were excluded. This threshold has been demonstrated in previous studies to significantly reduce the number of false positives [24,35].

### 6. Identification of viruses of emergence risk

We used the term ‘virus of emergence risk’ (VOER) is designate viruses that likely pose a greater threat of emerging in human populations (i.e., zoonotic virus), thereby distinguishing them from other, likely less impactful viruses. The identification of a virus as a VOER is based on three criteria: (i) phylogenetic clustering with established human pathogens or vector-borne virus groups, such as *Mammarenavirus, Norovirus, Orthornairovirus,* and *Morbillivirus*; (ii) that they had more than 80% amino acid identity in conserved genes (RdRp or DNA pol) to known pathogenic viruses of humans; and (iii) evidence of frequent co-species transmission, demonstrated by the presence of the virus in question in at least three host species from different mammalian families.

### 7. Virus distribution among organs within mammalian host species

We also quantified the total abundance (RPM) and detection frequency of each virus across three organs (gut, lung, and spleen) to evaluate their possible tissue preferences. For robustness, only sample groups where all three organ types had been sampled and sequenced were included (Supplementary Table 2). To enhance the robustness of our data, we excluded viruses detected in fewer than three libraries. The distribution data for each virus species across the organs were analyzed and visualized using the pheatmap package in R (version 4.1.1) [62]. To investigate potential cross-species virus transmission events, we utilized network visualization in R (version 4.1.1) using the igraph package [63].

### 8. Collection of data for ecological comparisons

To explore the impact of biodiversity and environmental factors on the diversity and composition of mammal-associated viromes, we gathered climate and land-use data for each sample site from public resources. Climate data were sourced from the TerraClimate data set [64], covering monthly climate variables—including six primary and eight derivative metrics—from 2021 to 2023. To reduce redundancy due to collinearity among climate variables, we conducted a principal component analysis. The first three principal components—CPC1, CPC2, and CPC3—accounted for 55.3%, 26.7%, and 6.3% of the total variance, respectively, cumulatively representing 88.3%. These components were subsequently utilized in all further statistical analyses to assess influences of climates on viral diversity and composition. Land-use data, obtained from the HYDE 3.2 database with a spatial resolution of 5 arcmin, were analyzed using the same method outlined previously [65]. The first two PCs, LPC1 and LPC2, explained 63.1% and 28.5% of the variance, respectively, totaling 91.6%. These components were then used to represent land-use patterns. Human population density data for 2022 were also sourced from the HYDE 3.2 database. Mammal richness data for each sampling site, aligned with the sampling year, were acquired from the International Union for Conservation of Nature mammal richness database. Monthly NDVI data from NASA’s MODIS MOD13A3 product for the years 2021 to 2023 were downloaded and averaged annually to calculate annual average NDVI values.

### 9. Statistic methods

All statistical analyses were conducted using R version 4.1.1 [62].

#### 9.1 Group selection

To minimize the confounding effects of uneven pooling sizes, we selected a subset of sample groups for our statistical analysis, focused on identifying the ecological drivers of viral diversity and cross-species transmission. We included sample groups from mammalian species with at least three replications (i.e., > 3 sample groups). We also restricted our analysis to groups containing lung, spleen, and gut, data as organ diversity can influence virus detection and composition. To account for environmental variability within the same group, we ensured that the standard deviation for principal component scores of climate (CPC1, CPC2) and land-use (LPC1, LPC2) variables remained below 1.2. We also verified that the median, mean, and variance of distances among sampling sites within a group did not exceed 50 km. This selection process resulted in 94 groups that satisfied these conditions (Supplementary Table 2), and all subsequent ecological analyses were performed based on this refined data set.

#### 9.2 Assessing viral richness and identifying key determinants

Consistent with previous studies [24,35], we used generalized linear models (GLMs) with negative binomial regression, as implemented in the MASS package [66] in R, to analyze the impact of host species, climate, and land-use on viral richness. The analysis incorporated data on host species, climatic variables (CPC1, CPC2, CPC3), and land use metrics (LPC1, LPC2, NDVI, log-transformed population density, mammal richness), in addition to average altitude and site count. For groups derived from multiple sampling sites, numerical values such as CPC1, LCP1, NDVI, log-transformed population density were averaged. Using the MuMIn package [67] in R, we systematically explored all possible variable combinations, selecting the most informative model based on the Akaike Information Criterion (AIC). This approach enabled a detailed assessment of each variable’s specific contribution, providing a nuanced understanding of their individual and combined effects on viral richness across mammalian hosts.

#### 9.3 Comparisons of virus transmission and identifying key determinants

Virome compositions across mammalian genera were visualized using t-SNE and differences were assessed through permutational multivariate analysis of variance (PERMANOVA), utilizing Jaccard distance calculated with the vegan package [68] in R. Viral genomic diversity within species was evaluated by aligning all sequence variants for each species using ClustalW and pairwise sequence identities were computed with the msa package [69] in R. We also explored the impact of host phylogenetic distance, climatic and land use variations, and spatial distance on the virus sharing patterns and viral genomic diversity using generalized linear models (GLM). The specific influences of each variable were quantified employing a systematic approach similar to the previously described model selection and assessment methodology.

## Acknowledgements

This work was supported by the National Natural Science Foundation of China (82341118, 32270160), Natural Science Foundation of Guangdong Province of China (2022A1515011854), Shenzhen Science and Technology Program (JCYJ20210324124414040 and KQTD20200820145822023), Major Project of Guangzhou National Laboratory (GZNL2023A01001), Guangdong Province “Pearl River Talent Plan” Innovation, Entrepreneurship Team Project (2019ZT08Y464), and the Fund of Shenzhen Key Laboratory (ZDSYS20220606100803007). Y. F. was supported by Yunnan Revitalization Talent Support Program Top Physician Project (XDYC-MY-2022-0074). Z.-H.G. was supported by National Natural Science Foundation of China (32360248 and 81660554) and by Support Plan for Talents in Yunnan (YNWR-MY-2018-035). E.C.H. was supported by an NHMRC (Australia) Investigator Award (GNT2017197) and by AIR@InnoHK administered by the Innovation and Technology Commission, Hong Kong Special Administrative Region, China.

We wish to thank the local Centers for Disease Control and Prevention in all 129 counties from 16 prefectures of Yunnan Province, for their assistance in specimen collection.

## Author Contributions

Conceptualization, Y.F., G.-D.L., E.C.H., Z.-H.G., and M.S.; Methodology, Y.F., G.-P.K., Y.-F.P., and M.S.; Investigation, Y.F., G.-P.K., Y.-F.P., J.W., W.-H.Y., W.-C.W., G.-D.L., and M.S.; Writing – Original Draft, G.-P.K., Y.-F.P., E.C.H., and M.S.; Writing – Review and Editing, All authors; Funding Acquisition, Y.F., Z.-H.G., and M.S.; Resources (sampling), Y.F., G.-P.K., W.-H.Y., H.P., J.W., X.H., L.-F.Y., and Z.-H.G.; Resources (Computational), G.-P.K., Y.-F.P., G.-Y.X., Y.-T.S., Q.-Y.G, X.L., and M.S.; Supervision, Y.F., D.-Y.G., G.-D.L., Z.-H.G., and M.S..

## Competing interests

The authors declare no competing interests.

## Supplementary information

**Figure S1.**
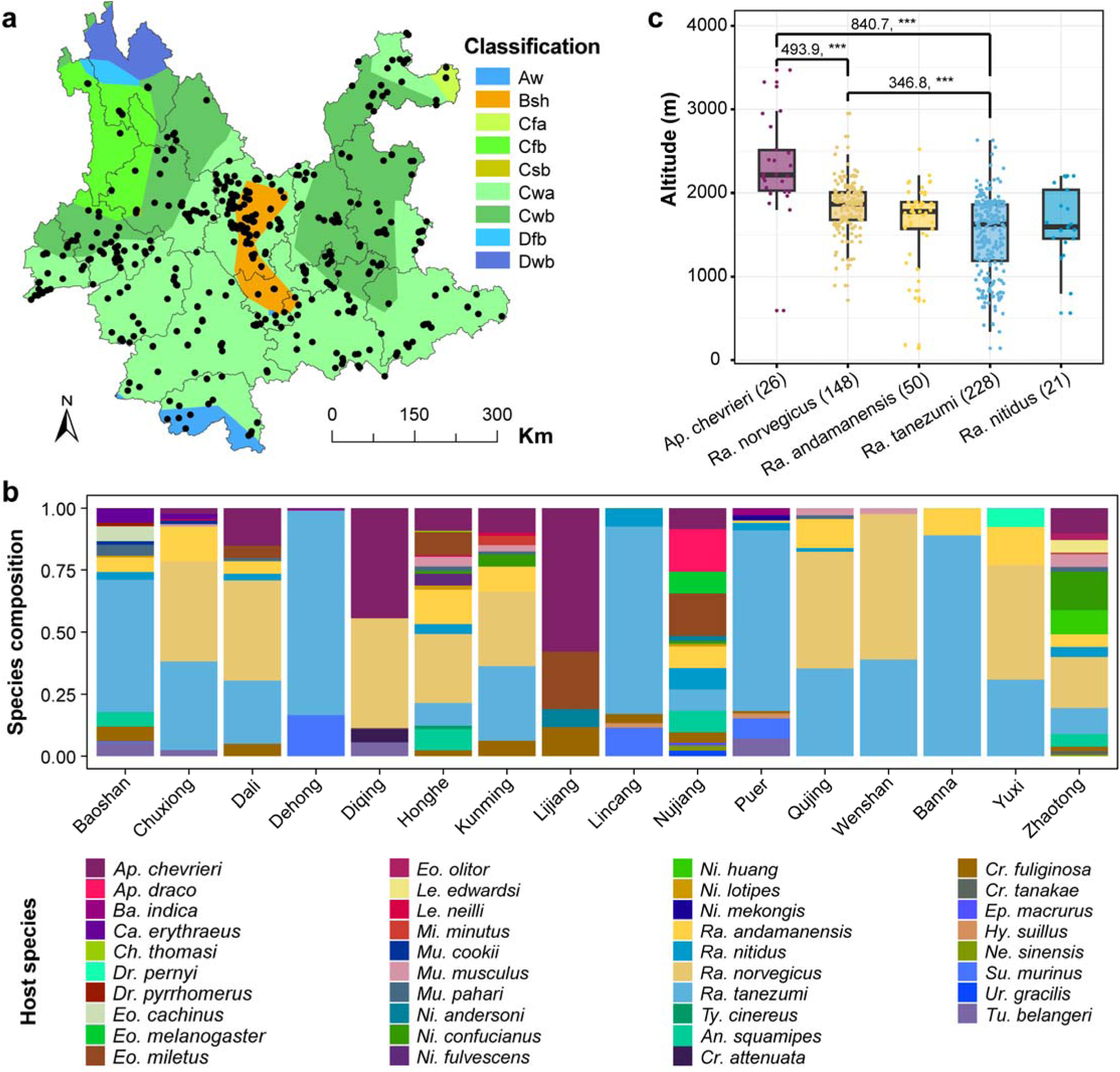
Climatic zones and mammalian species composition in this study. **(a)** Climatic zones within sampling area, classified according to the Köppen climate classification system, with black points indicating sampling sites. **(b)** Composition of mammalian species in samples collected across various prefectures. **(c)** Boxplot depicting the altitude distribution (meters) of specific rodent species, including only those five species collected from more than 20 different sites.

**Figure S2.**
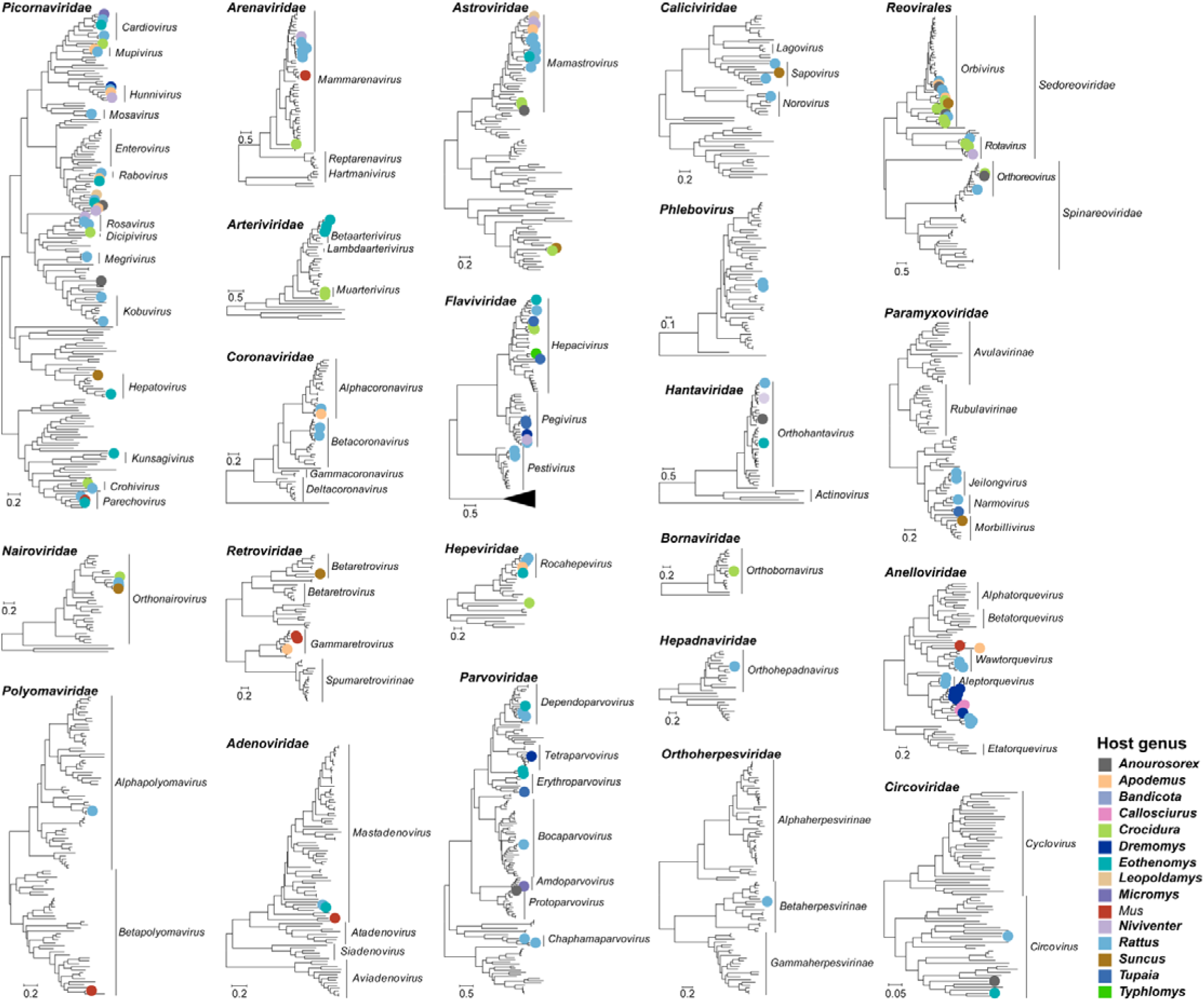
Phylogenetic diversity of mammalian viruses from 23 families identified in this study. Phylogenetic trees were estimated using a maximum likelihood method based on conserved protein sequences: RdRp for RNA viruses, reverse-transcriptase for the *Retroviridae*, the major capsid protein for the *Orthoherpesviridae*, LTAg for *Polyomaviridae*, ORF1 protein for the *Anelloviridae*, NS1 for the *Parvoviridae*, and DNA polymerase for other DNA viruses. Each tree is midpoint rooted with branch lengths scaled according to the number of substitutions per site. Viral species identified in this study are represented by dots, color-coded according to host genera, as detailed in the figure.

**Figure S3.**
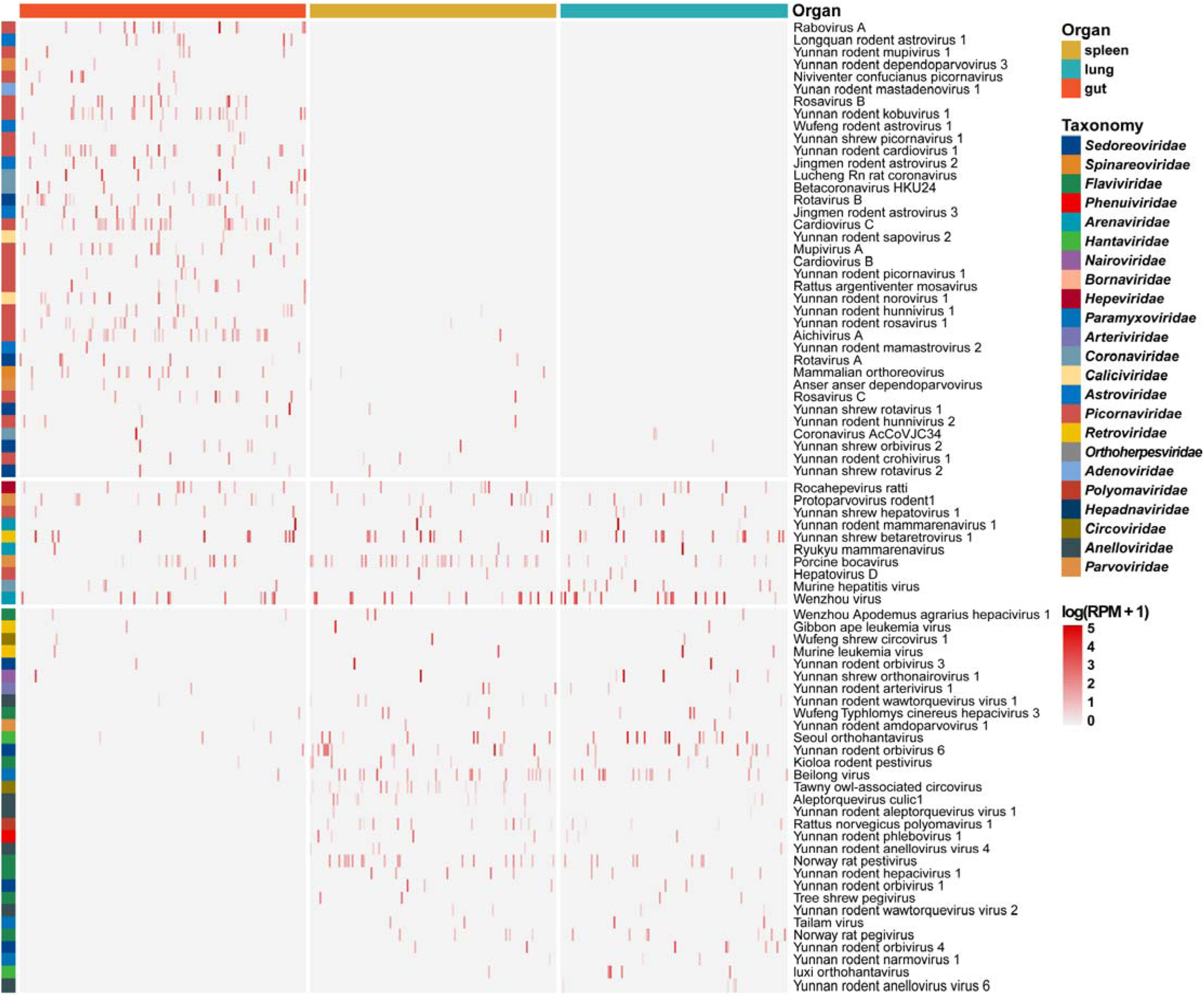
Distribution of virus abundance across different organs. Heatmap displays the abundance of each viral species across three organs (spleen, lung, and gut) in mammalian hosts examined in this study. The arrangement of libraries and viruses is organized by Canonical Correspondence Analysis (CCA).

**Figure S4.**
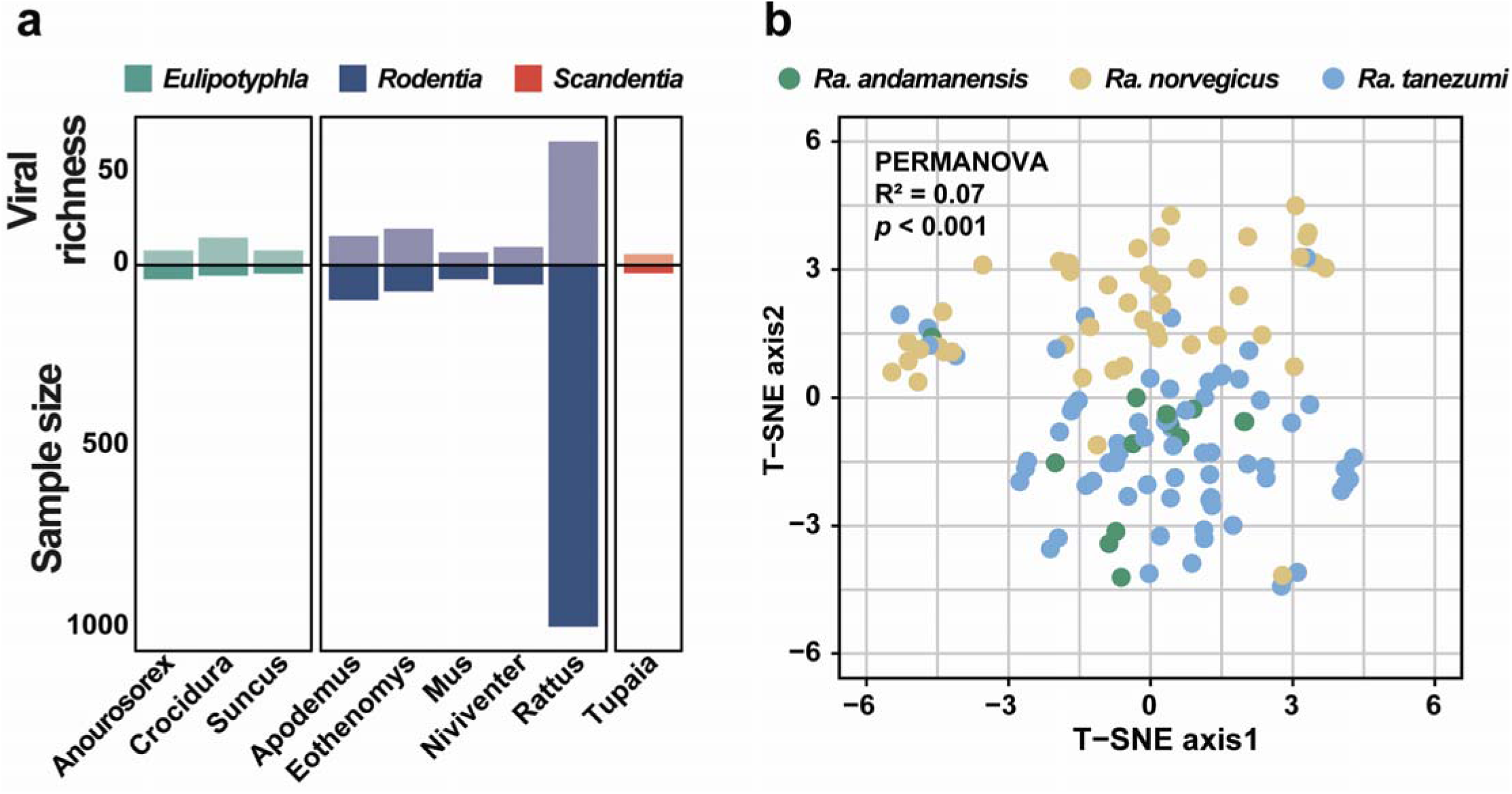
Viral richness and composition across host taxa. **(a)** Viral species identified within each genus of the orders Eulipotyphla, Rodentia, and Scandentia, including respective sample sizes. (**b)** t-SNE analysis showcasing the clustering of viral compositions among three rodent species within the genus Rattus: *Rattus tanezumi* (n = 67), *Rattus norvegicus* (n = 44), and *Rattus andamanensis* (n = 13). Statistical significance was assessed using PERMANOVA tests (two-sided) based on Jaccard distance with 1999 permutations.

**Figure S5.**
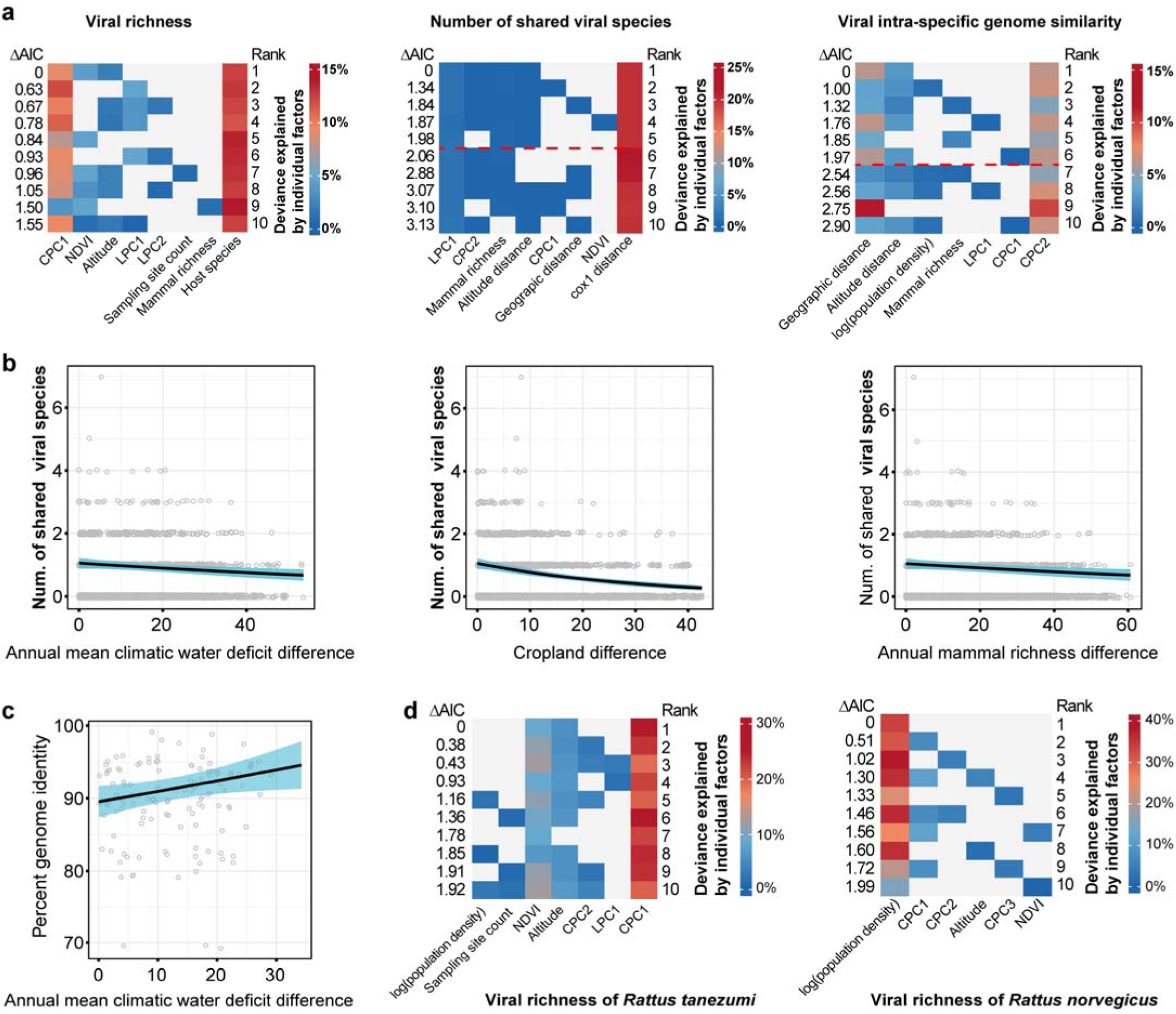
Environmental and host factors affecting viral species richness, composition, and intra-specific genomic diversity. **(a)** Relative effects of mammal species, climate, and land-use characteristics on viral richness, composition, and intra-specific genomic diversity. These effects are quantified by explained deviance in generalized linear models. The top 10 models selected by AIC are displayed, with the red dashed line indicating models significantly supported (ΔAIC <2). **(b)** The impact of differences in annual mean climatic water deficit, cropland presence, and mammal richness on viral sharing between pairs of sample groups. **(c)** The impact of differences in annual mean climatic water deficit on intra-specific viral genome similarity. **(d)** Relative effects of environmental characteristics on viral richness within groups of *Rattus tanezumi* and *Rattus norvegicus*.

**Supplementary Table 1.** Information on sampling sites and sample size of each mammal species.

**Supplementary Table 2.** Information of sample group and library in the current study.

**Supplementary Table 3.** Mammalian viruses identified in this study.

**Supplementary Table 4.** Viruses of emergence risk (VOERs) identified in this study.

## Notes

### Competing Interest Statement

The authors have declared no competing interest.

